# A non-catalytic role of RecBCD in Homology Directed Gap Repair and Translesion Synthesis

**DOI:** 10.1101/102509

**Authors:** Luisa Laureti, Lara Lee, Gaëlle Philippin, Vincent Pagès

## Abstract

The RecBCD complex is a key factor in DNA metabolism. This protein complex harbors a processive nuclease and two helicases activities that give it the ability to process duplex DNA ends. These enzymatic activities make RecBCD a major player in double strand break repair, conjugational recombination and degradation of linear DNA. In this work, we unravel a new role of the RecBCD complex in the processing of DNA single-strand gaps that are generated at DNA replication-blocking lesions. We show that independently of its nuclease or helicase activities, the entire RecBCD complex is required for recombinational repair of the gap and efficient translesion synthesis. Since none of the catalytic functions of RecBCD are required for those processes, we surmise that the complex acts as a structural element that stabilizes the blocked replication fork, allowing efficient DNA damage tolerance.

## INTRODUCTION

Genomes of all living organisms are constantly damaged by endogenous and exogenous stresses. Despite efficient repair mechanisms, some DNA lesions can escape repair and block the replicative polymerase. In order to bypass these “roadblocks” and complete replication, cells have developed two DNA Damage Tolerance (DDT) pathways identified both in prokaryotes and eukaryotes: 1) Translesion Synthesis (TLS), which employs specialized DNA polymerases able to replicate damaged DNA, with the potential to introduce mutations (1); 2) Damage Avoidance (DA) pathways (also named template switching), which use the information of the sister chromatid to bypass the lesion in a non-mutagenic way through homologous recombination mechanisms (2, 3). While the TLS pathway has been well characterized in the past few years, little is still known about Damage Avoidance pathways.

We have recently developed a genetic tool that enables us to monitor *in vivo* the exchange of genetic information between sister chromatids (*i.e*. DA events), following the insertion of a single lesion into the chromosome of *Escherichia coli* (4). We showed that after encountering a replication-blocking lesion, either on the lagging or the leading strand, the replication fork is able to restart downstream of the lesion leaving a single strand gap. Filling of this gap (also termed “single strand gap repair”, SSG repair) can be achieved either by TLS, or, to a higher frequency, by a DA mechanism that we named “Homology Directed Gap Repair” (HDGR). The HDGR pathway proved to be dependent on the bacterial recombinase RecA through the RecFOR pathway, already known to be involved in single strand gap repair (5-8). The major function of the RecFOR complex at a gapped DNA is to disassemble the filament of single strand binding protein (SSB) in order to load RecA and promote homologous recombination (9, 10). Interestingly, we also observed the participation of RecB in HDGR events (4, 11). RecB is part of the RecBCD complex, which is the key enzyme for initiation of recombinational repair of double-strand breaks (DSB) (12), conjugational recombination (13) and for degradation of linear DNA (also known as ExoV) in *E. coli* (5, 14). The RecBCD complex is composed of three distinct subunits (RecB, RecC and RecD) that together encompass several catalytic activities: DNA-dependent ATPase, DNA helicase, ssDNA endo and exonuclease, and dsDNA exonuclease. These activities enable RecBCD to be a potent and highly processive helicase and nuclease complex that processes duplex DNA ends and loads the recombinase RecA onto single-stranded DNA (ssDNA) during recombination events. The N-terminal regions of RecB and RecD contain a SF1 helicase motif (15), conferring a 3’5’ and 5’3’ helicase activity, respectively. This bipolar translocation is the basis for the characteristic velocity and processivity of RecBCD (it can unwind up to 30 kbp per binding event) (16, 17). The C-terminal region of RecB contains the nuclease domain as well as the RecA interaction domain. A specific DNA sequence named Chi (5’-GCTGGTGG-3’) regulates all the catalytic activities of RecBCD (reviewed in (5, 14)). The function of RecC still needs to be completely elucidated, however *recC* mutants seem to point towards a role in Chi recognition (18, 19).

Traditionally, the RecBCD complex has always been associated with the repair of double-strand breaks, while the RecFOR pathway was associated with the repair of single strand gaps formed upon replication fork stalling followed by re-priming downstream of the lesion (5, 6). However, it has been shown that when part (or all) of the enzymatic machinery of RecBCD is affected, RecQ helicase and RecJ nuclease (that are part of the RecFOR pathway) can achieve the resection of DNA ends, while RecFOR loads RecA nucleofilament, allowing the cell to be completely proficient for DSB repair (20, 21). In contrast, until our recent study, no evidence pointed towards a role for RecBCD in SSG repair. Indeed, we showed that inactivation of the *recB* gene leads to a decrease in HDGR events, even in the presence of a functional RecFOR pathway, suggesting that RecBCD does not act as a backup, but has its own contribution. To perform an efficient HDGR mechanism, both RecBCD and RecFOR complexes are necessary since the double mutant showed a phenotype similar to that of a *recA* mutant (i.e. an almost complete abolition of HDGR events). In the present work, we are further elucidating the role of RecBCD in the HDGR pathway. We demonstrate that the RecBCD complex is involved in SSG repair in a non-canonical way that is distinct from its DSB repair functions. Indeed, none of the characteristic enzymatic activities of RecBCD (*i.e*. nuclease, helicase and RecA-loading) are required for its participation to HDGR mechanisms. Furthermore, we find the TLS pathway to be strongly affected in the absence of RecBCD. We suggest that the RecBCD complex plays an unprecedented structural role in single strand gap repair that is necessary for both HDGR and TLS pathways.

## MATERIALS AND METHODS

### Bacterial strains and growth conditions

All *E. coli* strains used in this work are derivative of strains FBG151 and FBG152 (22, 23) and are listed in Supplementary Table 1. Strains were grown on solid and in liquid Lysogeny Broth (LB) medium. Gene disruptions of *recA, recF, recO, recB, recD, sulA, mutS* and *uvrA* were achieved by the one-step PCR method (24). To obtain the *recD^K177Q^* strain, the *recD* gene has been first cloned into the pKD4 vector (24) digested by *Nde*I and then we performed site-specific mutagenesis to obtain the K177Q substitution. We amplified by PCR the *recD^K177Q^* allele together with the kanamycin resistance gene and performed the one-step PCR method to obtain the *recD^K177Q^* strain. Strain RIK174 that contains the *recB^D1080A^* allele (25) was obtained from the Gene Stock Center. In order to transduce this allele in our strains, we inserted by the one-step PCR method a kanamycin resistance gene cassette in the intergenic region *ppdA-thyA* 0.25 min away from the *recB* gene. The presence of point mutations in the strains EVP629, 630, 654, 655, 658, 659, 712, 713 has been verified by sequencing. All strains carry a plasmid that allows the expression of the int–xis genes under the control of IPTG. Following the site-specific recombination reaction, the lesion is located either in the lagging strand (FBG151 derived strains) or in the leading strand (FBG152 derived strains). Antibiotics were used at the following concentrations: ampicillin 50 or 100 μg/ ml; tetracycline 10 μg/ml, kanamycin 100 μg/ml, chloramphenicol 30 μg/ml. When necessary IPTG and X-Gal were added to the medium at 0.2mM and 80 μg/ml, respectively.

### Plasmids

A list of all the plasmids used is this study is provided in Supplementary Table 2.

pVP135 expresses the integrase and excisionase (int–xis) genes from phage lambda under the control of a *trc* promoter that has been weakened by mutations in the -35 and the -10 region (26). Transcription from Ptrc is regulated by the *lac* repressor, supplied by a copy of *lacI^q^* on the plasmid. The vector has been modified as previously described (22). pKN13 is similar to pVP135 except that it possesses a chloramphenicol resistance gene instead of a kanamycin resistance gene.

pLL58 and pLL59 are derived from pKN13 and contain the *recA* gene or the *recA730* allele, respectively, in continuity with the *xis-int* operon. The genes together with their ribosome-binding site have been cloned in pKN13, previously digested by HindIII and blunt ended.

pVP146 is derived from pACYC184 plasmid where the chloramphenicol resistance gene has been deleted by BsaAI digestion and re-ligation. This vector, which carries only the tetracycline resistance gene, serves as an internal control for transformation efficiency.

pVP141-144, pGP1, 2 and 9 are derived from pLDR9-attL-*lacZ* as described in (22). pLL1 and pLL2c are derived from pVP141 and contain several genetic markers as previously described (4). All these plasmid vectors contain the following characteristics: the ampicillin resistance gene, the R6K replication origin that allows plasmid replication only if the recipient strain carries the *pir* gene (27), and the 5’ end of the *lacZ* gene in fusion with the attL site-specific recombination site of phage lambda. The P’3 site of attL has been mutated (AATCATTAT to AAT<u>TA</u>TTAT) to avoid the excision of the plasmid once integrated (28). These plasmids are produced in strain EC100D pir-116 (from Epicentre Biotechnologies, cat# EC6P0950H) in which the *pir-116* allele supports higher copy number of R6K origin plasmids. Vectors carrying a single lesion for integration were constructed as described previously (22) following the gap-duplex method (29). A 13-mer oligonucleotide, 5'-GCAAG<u>TT</u>AACACG-3', containing no lesion or a TT(6-4) lesion (underlined) in the HincII site was inserted either into the gapped-duplex pLL1/2c leading to an out of frame *lacZ* gene (to measure HDGR) or into the gapped-duplex pGP1/2 leading to an in frame *lacZ* gene (to measure TLS). A 15-mer oligonucleotide 5'-ATCACCGGC<u>G</u>CCACA-3’ containing or not a single G-AAF adduct (underlined) in the NarI site was inserted into the gapped-duplexes pVP141-142 or pVP143-144 to score respectively for TLS0 Pol V-dependent and for TLS-2 Pol II-dependent. A 13 - mer oligonucleotide, 5’-GAAGACCT<u>G</u>CAGG, containing no lesion or a dG-BaP(-) lesion (underlined) was inserted into the gapped-duplex pVP143/pGP9 leading to an in frame *lacZ* gene (to measure TLS).

### Monitoring HDGR and TLS

To 40 μL aliquot of competent cells, prepared as previously described (22), 1 ng of the lesion-carrying vector mixed with 1 ng of the internal standard (pVP146) was added and electroporated in a GenePulser Xcell from BioRad (2.5 kV, 25 μF, 200 Ω). Cells were first resuspended in super optimal broth with catabolic repressor (SOC), then diluted in LB containing 0,2 mM IPTG. Cells were incubated for 45 min at 37 °C. Part of the cells were plated on LB + 10 μg/mL tetracycline to measure the transformation efficiency of plasmid pVP146, and the rest were plated on LB + 50 μg/mL ampicillin + 80 μg/mL X-gal to select for integrants (Amp^R^) and to visualize HDGR or TLS events (*lacZ+* phenotype depending on the vector used). Cells were diluted and plated using the automatic serial diluter and plater EasySpiral Dilute (Interscience). Colonies were counted using the Scan 1200 automatic colony counter (Interscience). Integration rate, transformation efficiency and plating efficiency vary according to the genetic backgrounds. The integration rate is in the range of 2,000 clones per picogram of vector for our parental strain. For *rec-* strains whose viability is affected and plating efficiency reduced, this rate can drop to ∼200 clones per picogram.

We plated before the first cell division; therefore, following the integration of the pLL1/2c vector, sectored blue/white colonies represent HDGR events; sectored pale blue/white colonies represent TLS events and pure white colonies represent damaged chromatid loss event (Figure 1). Following integration of the pVP141/142, pVP143/144, pGP1/2, pVP143/pGP9 vectors, sectored blue/white colonies represent TLS events. The relative integration efficiencies of lesion-carrying vectors compared with their lesion-free homologues, normalized by the transformation efficiency of pVP146 plasmid in the same electroporation experiment, allow the overall rate of lesion tolerance to be measured.

**Figure 1.**
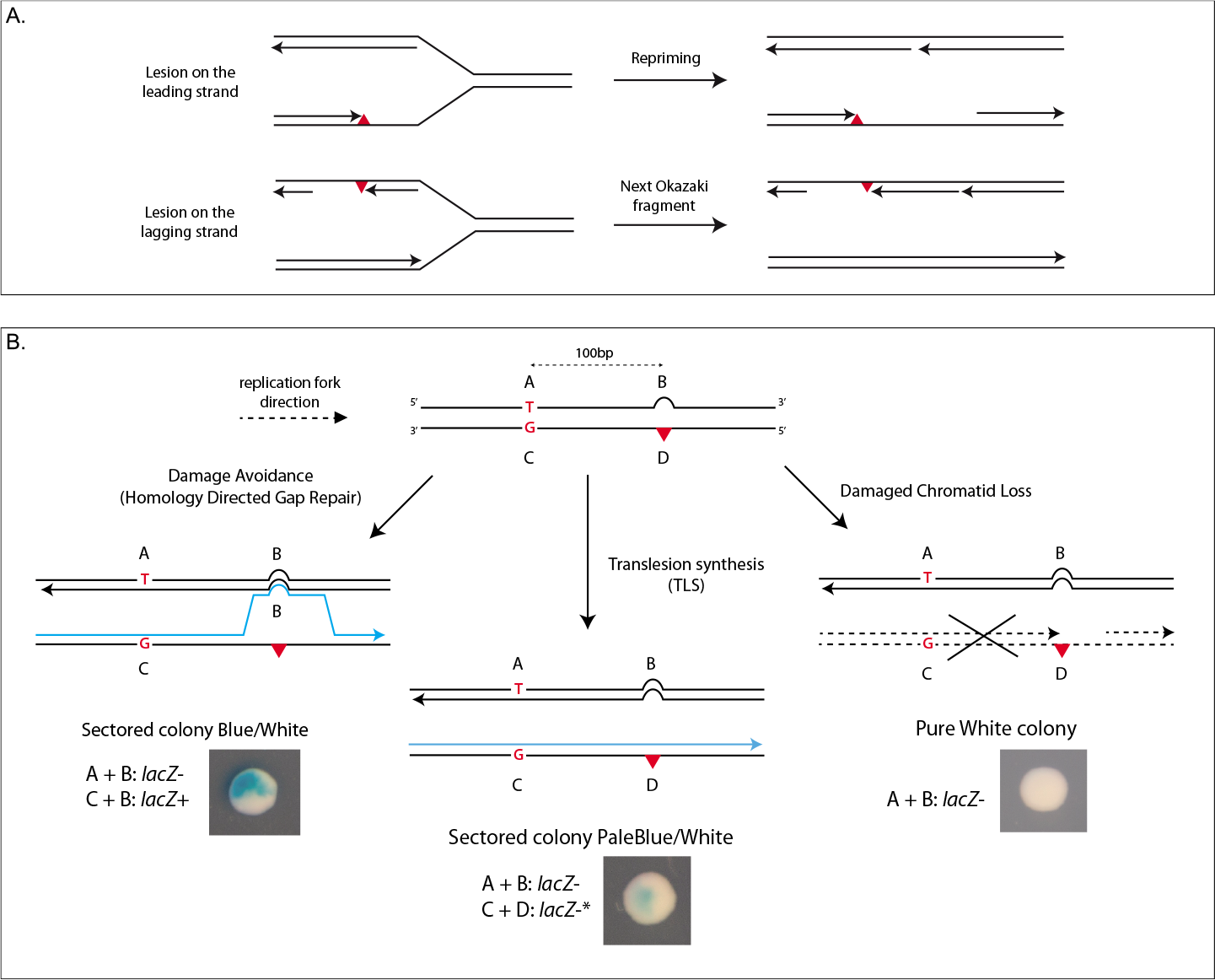
A. Model of the replication fork encountering a lesion in the leading or lagging strand. B. Genetic system to monitor sister-strand exchange mechanisms (modified from (4)). The scheme represents the situation in which the lesion (red triangle) is located in the 5’′end of the *lacZ* gene in the leading strand. The damaged strand containing the marker D, where the lesion is located, and the marker C, placed 100 bp upstream the lesion, contains a +2 frameshift in order to inactivate the *lacZ* gene. Opposite to the lesion we introduced a +4 loop (marker B) that restores the reading frame of *lacZ*, and in the same strand we added marker A that contains a stop codon. Therefore the two strands are *lacZ*-. Only a mechanism of HDGR by which replication has been initiated on the damaged strand (incorporation of marker C), and where a template switch occurred at the lesion site (leading to incorporation of marker B) will restore a *lacZ+* gene (the combination of markers C and B contains neither a stop codon nor a frameshift). Using the same system we can also score for TLS events (combination of marker C and D), as sectored pale blue/white colonies, and for damaged chromatid loss events (combination of marker A+B), as pure white colonies. *For the combination of marker C and D we observed a leaky activity of the -galactosidase due to a translational frameshift.

## RESULTS

### The nuclease domain of RecB is not required for HDGR mechanism

In order to assess the role played by the different subunits of RecBCD in HDGR, we used a genetic system that we previously developed (4). Briefly, a vector containing a single replication-blocking lesion is integrated in the bacterial chromosome by mean of the phage lambda integrase. The vector carries a combination of four genetic markers (Figure 1) that allows to directly visualize the exchange of genetic information between the damaged strand and the non-damaged sister chromatid. Using this as say, we previously showed that sister-strand exchange events (named Homology Directed Gap Repair) are the major DDT pathway. When cells fail to fill the ssDNA gap, they can still survive by replicating their non-damage d chromatid and losing the damaged one (4). We named the seevents “damaged chromatid loss”. We also showed that the HDGR path way is dependent on the recombinase RecA, mainly through the RecFOR pathway and to a lesser extent through the action of RecB (Figure 2, and see (4)). The deletion of *recB* gene was indeed accompanied by a decrease in HDGR events of ∼30% when the lesion is located on the lagging strand, and ∼15% when on the leading strand (Figure 2 and (4)). When the HDGR pathway is affected, it can be compensated by an increase in damaged chromatid loss events, as we can clearly see in a *recF-* strain and to a lesser extent in the *recB-* strain (Figure 2). In some cases, a decrease in HDGR can also be accompanied by a decrease in survival, as in the *recB-* strain for the leading strand or in the *recB recF* double mutant (Figure 2). A loss of survival is attributed to the presence of unrepaired lesions on the opposite strand that prevents damaged chromatid loss to occur and leads to lethality (4). Since it is the first time that the RecBCD complex appears to be involved in single strand gap repair, we undertook to further explore its role in the HDGR mechanism.

**Figure 2.**
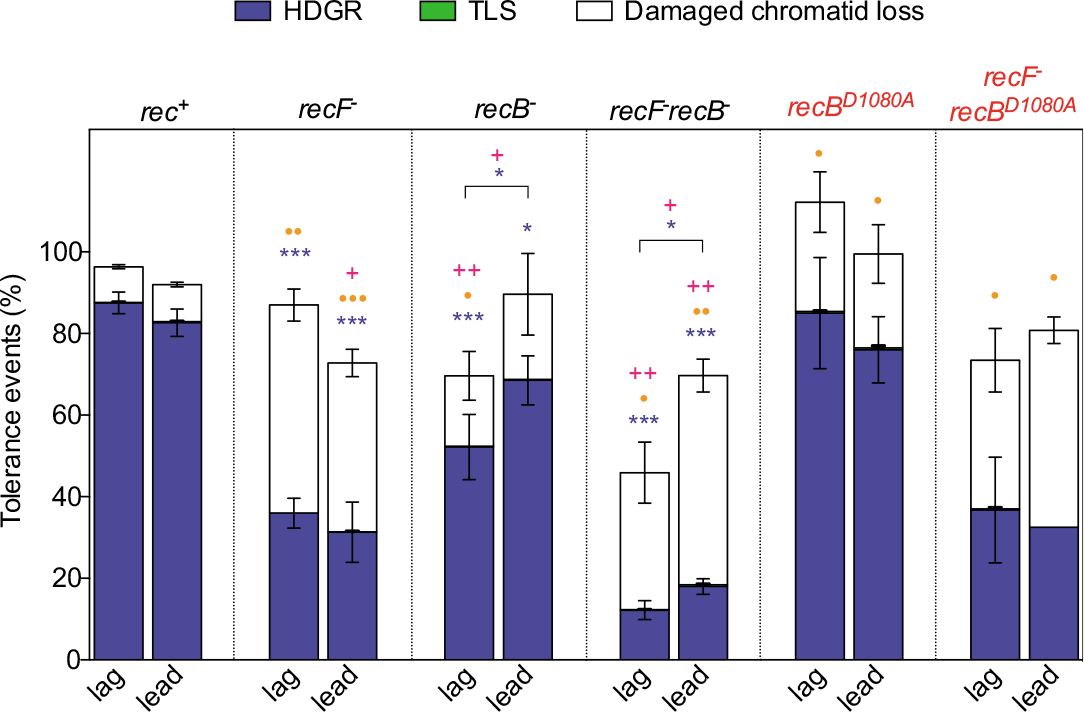
Partitioning of DDT pathways in the *recB* nuclease deficient strains (*recB^D1080A^*). The graph represents the partition of DDT pathways in the presence of the UV lesion TT(6-4) inserted in different recombinant deficient strains. The lesion has been inserted in either the leading (lead) or the lagging (lag) strands of *E. coli* chromosome. Tolerance events (Y axis) represent the percentage of cells able to survive in presence of the integrated lesion compared to the lesion-free control. *rec+* corresponds to our parental strain, recombination proficient. The data for *rec+, recF-, recB- and recF- recB-* strains have been previously published (4). *recB^D1080A^* strains deficient for the nuclease activity are indicated in red. The data represent the average and standard deviation of at least three independent experiments. T-test was performed to compare values from the different mutants to the *rec+* strain, with the exception of *recF-recB^D1080A^* whose values have been compared to the single *recF* mutant. We also compare values for the leading and lagging orientation of the *recB* and *recF recB* mutant strains. For HDGR: \**P* < 0.05; \*\**P* < 0.005; \*\*\**P* < 0.0005. For Damaged chromatid loss: •*P* < 0.05; ••*P* < 0.005; •••*P* < 0.0005. For survival: +*P* < 0.05; ++*P* < 0.005.

In the present study, all experiments were conducted in a parental strain where mismatch repair (*mutS*) has been inactivated (to prevent corrections of the genetic markers), as well as nucleotide excision repair (*uvrA*), to avoid excision of the lesion and to focus on lesion tolerance events. To measure HDGR events, we used the UV-induced thymine-thymine pyrimidine(6-4) pyrimidone photoproduct [TT(6-4)] blocking lesion. To measure TLS events, we also employed two known guanine adducts, the N - 2 - acetylaminofluorene (G-AAF) and the benzo(a)pyrene (dG-BaP(-)).

RecB is the major subunit of RecBCD complex and contains the nuclease domain, one of the two helicase domains and the domain for RecA interaction. Acting as a processive nuclease, RecB is able to degrade both strands of a blunt double strand DNA (dsDNA) template, with a preference for the 3’-end. In order to investigate a possible role of the nuclease domain of RecB in HDGR, we used the previously characterized *recB^D1080A^* nuclease dead allele (30) that contains a single point mutation in the catalytic core of the nuclease domain that prevents Mg^2+^ binding. The integration of our lesion-containing vector into the *recB^D1080A^* strain showed a level of HDGR similar to the parental strain (Figure 2), indicating that the nuclease activity of RecB is dispensable for HDGR. Noteworthy, the bias lagging *vs* leading in the HDGR level observed in the *recB* deficient strain is not observed in the nuclease dead allele.

*In vivo* studies showed that the *recB^D1080A^* strain is still proficient in DSB repair but entirely relies on the RecFOR complex for this (31, 32). Indeed, the D1080A mutation seems to also alter the RecA loading capacity of the RecBCD complex (33), but this is compensated *in vivo* by the RecFOR complex, the other mediator of RecA loading. We previously showed that a *recF recB* double mutant strain presents a strong defect in HDGR similar to a *recA* deficient strain, which suggested independent roles for both RecF and RecB in HDGR. In contrast, as shown in Figure 2, the *recF recB^D1080A^* double mutant is no more deficient in HDGR than the *recF* single mutant. This result shows that the contribution of RecBCD to HDGR does not involve its RecA loading activity, nor the processing of DSB that would have arisen from the single lesion.

Since the RecOR complex has been shown to be able to load RecA even in the absence of RecF (10, 21, 34, 35), we tested whether RecOR was able to compensate the RecA loading defect in the *recB^D1080A^* strain. As shown in Supplementary Figure 1, the defect in HDGR in a *recO-* strain is similar to the one in the *recF-* strain, and the *recO recF* double mutant doesn’t show any additional defect proving that the two genes are epistatic. More importantly, a *recO recB^D1080A^* double mutant behaves like a *recO* single mutant confirming the result obtained in the *recF recB^D1080A^* strain, and the above conclusion.

### The whole RecBCD complex participates to HDGR

Next, we wanted to address the question of whether RecD was dispensable for the HDGR activity of the complex. *In vivo* RecB needs at least RecC to be functionally active and mutants in either *recB* or *recC* gene show a similar phenotype, *i.e*. recombination deficiency, increased sensitivity to DNA damaging agents and decrease in cell viability (5). On the contrary, a *recD* null mutant even though deficient for DNA degradation (36) is proficient in recombination and DNA repair because the RecBC complex still possesses the 3’5’ helicase activity of RecB to unwind dsDNA and the ability to load RecA (37, 38). Since the nuclease activity of RecB turns out to be dispensable for HDGR and that RecD is the subunit that controls the nuclease activity of the complex, one would expect the deletion of *recD* not to affect HDGR level (as in a *recB^D1080A^* strain). Unexpectedly, following the integration of our lesion-containing construct in a *recD* deficient strain, we observed a decrease in HDGR similar to a *recB* deficient strain. However, while in a *recB-* strain HDGR and survival are more affected when the lesion is located on the lagging strand, in a *recD-* strain HDGR and survival decrease more significantly when the lesion is located on the leading strand (Figure 3). This result indicates that the RecD subunit, together with the rest of the complex, is required for HDGR; and that the helicase activity of RecB is dispensable since it is functional in a *recD* deficient strain and yet, we observe a decrease in the level of HDGR

**Figure 3.**
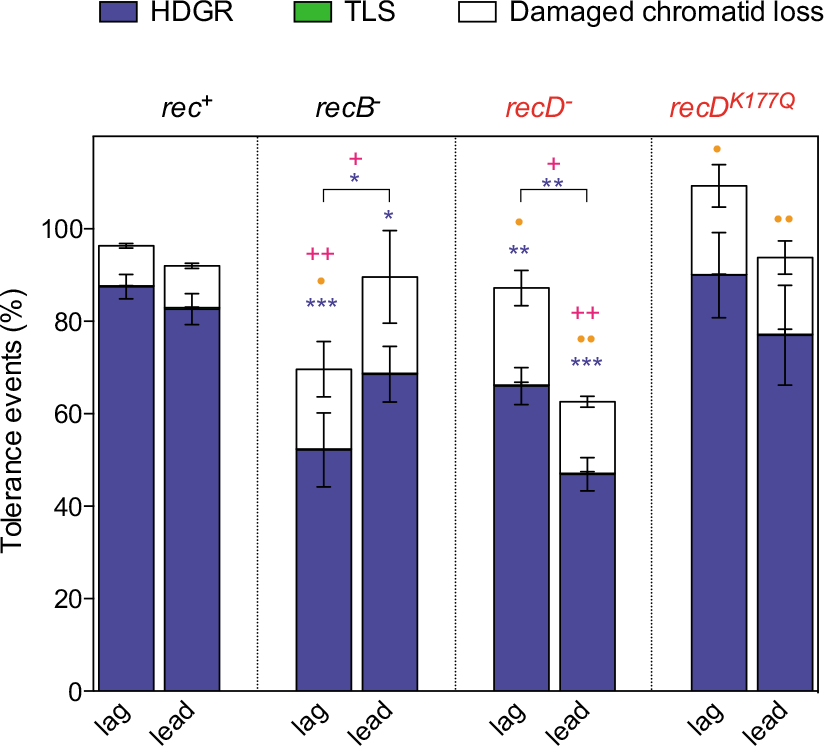
Partitioning of DDT pathways in *recD* mutant strains. The graph represents the partition of DDT pathways in the presence of the UV lesion TT(6-4) inserted in the *recD*– and the *recD^K177Q^* mutant strains (indicated in red). The lesion has been inserted in either the leading (lead) or the lagging (lag) strands of *E. coli* chromosome. Tolerance events (Y axis) represent the percentage of cells able to survive in presence of the integrated lesion compared to the lesion-free control. *rec+* corresponds to our parental strain, recombination proficient. The data for *rec+* and *recB–* strains have been previously published (4). The data represent the average and standard deviation of at least three independent experiments. T-test was performed to compare values from the different mutants to the *rec+* strain. We also compare values for the leading and lagging orientation of the *recB* and *recD* mutant. For HDGR: \**P* < 0.05; \*\**P* < 0.005; \*\*\**P* < 0.0005. For Damaged chromatid loss: •*P* < 0.05; ••*P* < 0.005; •••*P* < 0.0005. For survival: +*P* < 0.05; ++*P* < 0.005.

The RecD subunit not only controls the nuclease activity of the complex, but it also harbors its own 5’3’ DNA helicase activity. The decrease in HDGR level observed in the *recD* deficient strain could be due to the absence of this helicase activity. Therefore, to assess whether the RecD 5’3’ helicase activity is needed for HDGR, we constructed a helicase dead *recD^K177Q^* mutant where the Lys177Gln substitution in the Walker A motif of the ATPase domain of RecD is known to inactivate the helicase activity (17). Following the insertion of the TT6-4 lesion construct in the *rec^DK177Q^* strain, we did not observe any decrease in HDGR in contrast to the reduced HDGR levels measured in the *recD* deficient strain (Figure 3). This shows that the 5’3’ helicase activity of RecD subunit is also not required for HDGR. Altogether, these results clearly indicate that none of the helicases activities of RecBCD are required, but rather that the entire RecBCD complex participates in the HDGR mechanism.

### RecBCD complex is not a mediator of RecA loading at a single strand gap

Our data indicate that the RecBCD complex is involved in the HDGR mechanism, independently of its nuclease or helicase activities, contrarily to its role played in DSB repair. As previously mentioned RecBCD is, together with RecFOR, a mediator of RecA loading onto ssDNA. While the mediator activity of RecBCD is classically associated with its nuclease and helicase activities, we raised the question whether RecBCD could act as a mediator of RecA during SSG repair, without involving its helicases and nucleases activity. To address this question, we used a specific allele of RecA, the *recA730* (E38K) allele, that is able to load itself onto ssDNA without the help of its mediators (39). Previous studies demonstrated *in vivo* and *in vitro* that this allele partially complements the phenotype associated with a *recF(OR)* deficient strain (21, 40). We modified our plasmid expressing the lambda excisionase/ integrase under an inducible promoter (22) by adding either the *recA730* allele (pLL59) or the wild-type copy of the *recA* gene (pLL58) in order to express these alleles in the recipient strains. After ensuring that both plasmids were able to complement a *recA* deficient strain (Supplementary Figure 2), we transformed those plasmids in a *recF* deficient strain and monitored HDGR levels. As expected, despite the absence of the RecA mediator (RecF) the level of HDGR is significantly increased (by ∼2 folds) upon expression of the *recA730* allele, while no increase is observed upon expression of the wild-type *recA* gene (Figure 4). The *recA730* allele doesn’t fully complement the *recF* defect, as previously shown (21, 40) and as expected since it also fails to fully complement the *recA*- strain (Supplementary Figure 2). This result confirms that the defect in HDGR observed in the *recF*- strain is due to a defect in the RecA loading activity. Next, we transformed in the *recB* deficient strain the plasmids containing either the *recA730* allele or the wild-type *recA* gene. No increase in HDGR level was observed when the *recA730* allele was expressed (Figure 4), suggesting that the role of RecBCD complex in lesion tolerance is not to mediate the loading of RecA on the single strand gap.

**Figure 4.**
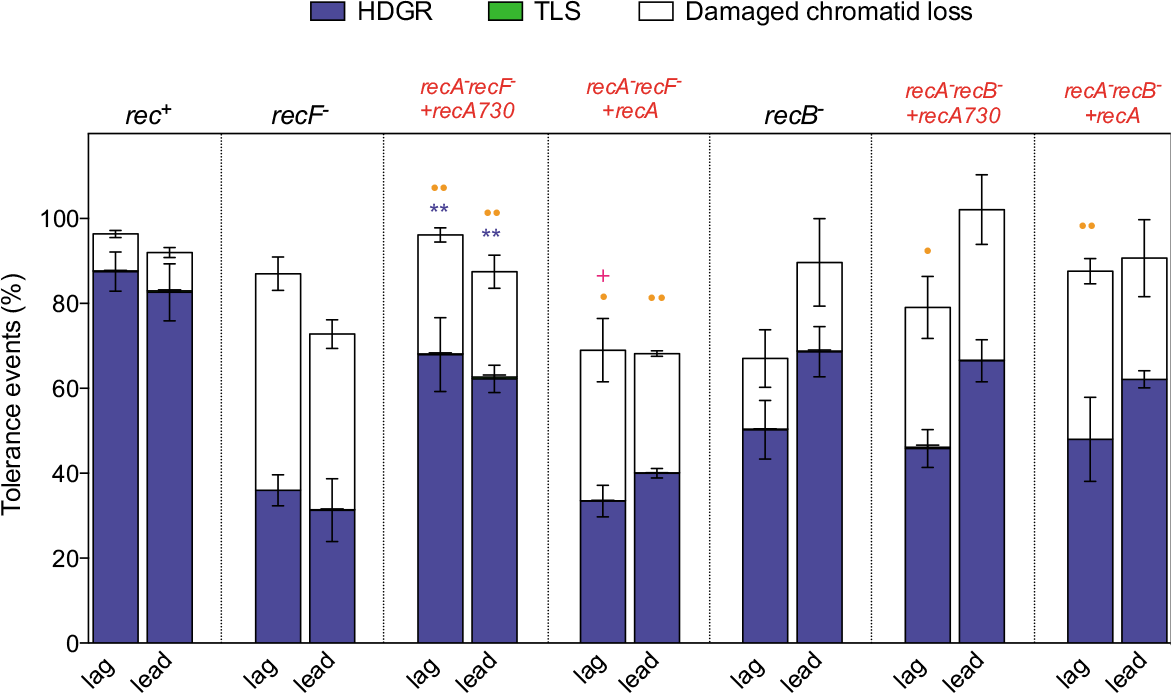
Partitioning of DDT pathways in strains expressing the *recA730* allele. The graph represents the partition of DDT pathways in the presence of the UV lesion TT(6-4) in strains expressing the *recA730* allele or the wild-type copy of RecA (indicated in red). In those strains the *sulA* gene has been also inactivated to avoid cell division blockage because of the constitutively SOS activation due to the *recA730* allele. The plasmid pLL59 contains the *recA730* allele, while the plasmid pLL58 contains the wild-type copy of *recA* gene. The lesion has been inserted in either the leading (lead) or the lagging (lag) strands of *E. coli* chromosome. Tolerance events (Y axis) represent the percentage of cells able to survive in presence of the integrated lesion compared to the lesion-free control. *rec+* corresponds to our parental strain, recombination proficient. The data for *rec+, recF*- and *recB-* strains have been previously published (4). The data represent the average and standard deviation of at least three independent experiments. T-test was performed to compare values from the double mutants (*recA recF* or *recA recB*) to the their correspondent single mutant strain (*recF* or *recB*). For HDGR: \**P* < 0.05; \*\**P* < 0.005. For Damaged chromatid loss: •*P* < 0.05; ••*P* < 0.005.

### The RecBCD complex is also involved in the TLS pathway

Since none of the known catalytic activities of RecBCD seems to be required for its role in HDGR, we hypothesized that RecBCD could play a structural role, possibly by stabilizing or helping in the stabilization of the stalled replication fork. If such stabilization is necessary for an efficient bypass of the lesion, we reasoned that it would be required not only for the HDGR mechanism, but also for the TLS pathway. Since TLS events at the TT(6-4) lesion are very rare events (≤ 0.5%) (Figure 5 and (22)), it is difficult to measure a significant decrease in the *recB* deficient strain (Figure 5). For this reason, we monitored the effect on TLS at two other lesions: the guanine adducts N-2-acetylaminofluorene (G-AAF) and benzo(a)pyrene (dG-BaP(-)) that show a higher basal level of TLS in the parental strain (see Figure 5 and (41)). When the G-AAF lesion is introduced in the *Nar*I sequence, a potent mutation hotspot, TLS can be mediated either by Pol V (TLS0, non-mutagenic) or by Pol II (TLS-2, frameshift -2) (42). The dG-BaP(-) lesion is bypassed by Pol IV (TLS0, non-mutagenic) (43, 44). We used our previously described integration assay that allows to specifically monitor TLS events (22, 41): in a *recB* deficient strain, we observed a substantial decrease in the bypass mediated by all three TLS polymerases (Figure 5). To ensure that this effect was not due to the catalytic activities of the RecBCD complex, we also measured TLS in the nuclease and helicase dead mutants, *recB^D1080A^* and *recD^K177Q^*: we didn’t observe any decrease in the TLS level in these mutants (Figure 5). However, both *recB^D1080A^* and *recD^K177Q^* strains are still proficient in DSB repair. To exclude the potential involvement of the formation and repair of a DSB that would be involved in the TLS pathway, we measured TLS in a *recF recB^D1080A^* and in a *recO recB^D1080A^* strains that are known to be impaired for DSB repair (31). We didn’t observe any diminution in the level of TLS in these strains when compared to the correspondent single mutant *recF* and *recO* (Supplementary Figure 3). It appears therefore that in addition to its role in HDGR, RecBCD is also required for an efficient TLS pathway independently of its catalytic activities and of its capacity to repair DSB.

**Figure 5.**
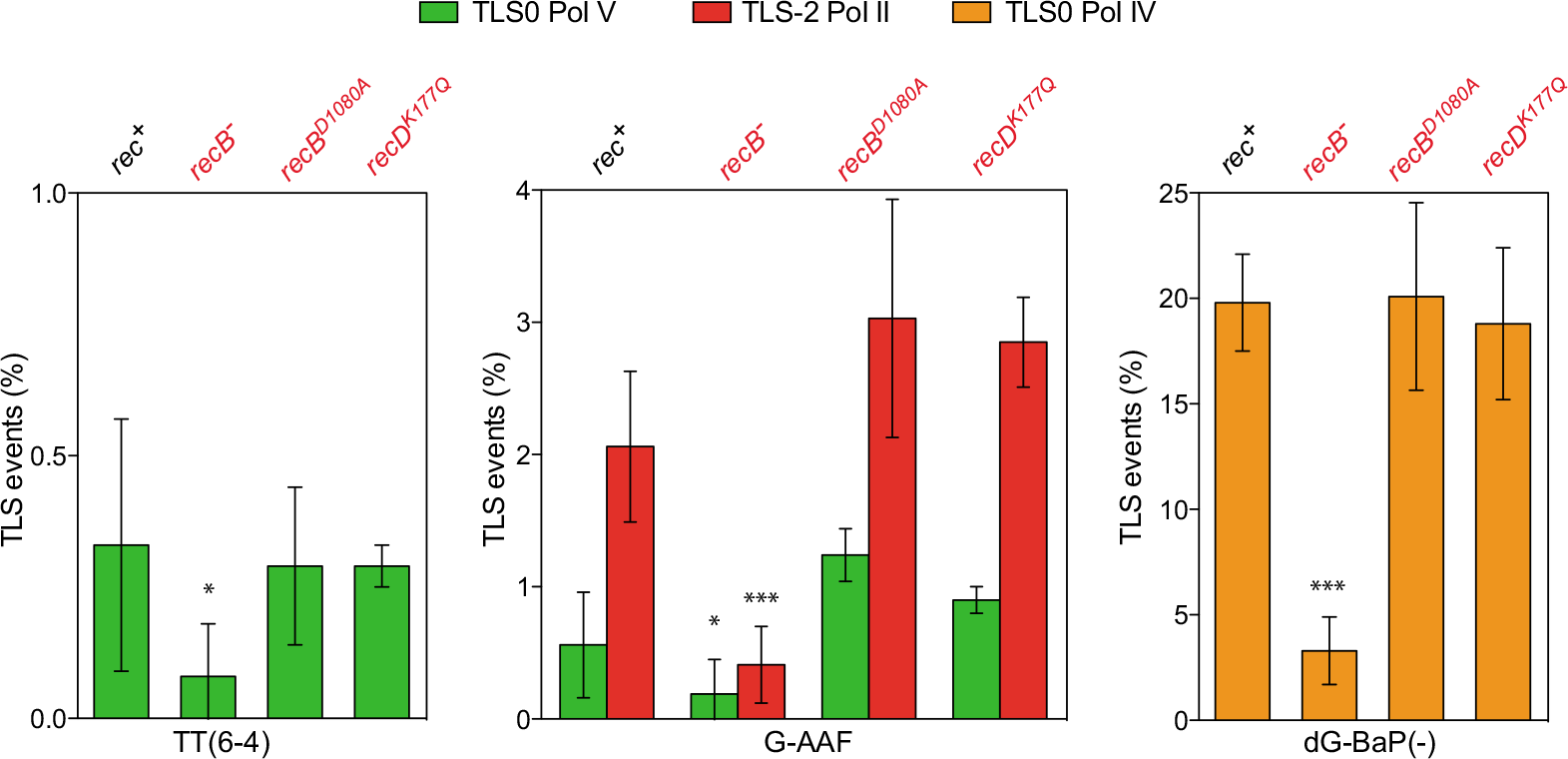
RecBCD complex is involved in TLS pathway. The graph represents the percentage of TLS events in the presence of three different replication blocking lesions (TT6-4, G-AAF, dG-BaP(-)) in the *recB-, recB^D1080A^* and *recD^K177Q^* strains (indicated in red) in comparison with a *rec+* strain. The data for *rec+* strain have been previously published (41, 49). The data represent the average and standard deviation of at least three independent experiments in which the lesion has been inserted either in the leading or the lagging strand. The data from leading and lagging strands have been pooled together because no significant difference was observed between the two orientations. T-test was performed to compare TLS values from the *recB-* strain to the *rec+* strain. \**P* < 0.05; \*\**P* < 0.005; \*\*\**P* < 0.0005.

## DISCUSSION

The present work aims at elucidating the role of RecBCD in DNA damage tolerance. RecBCD is one of the most fascinating and studied multienzymatic complexes in bacteria. It is the major recombinational pathway in *E. coli* responsible for the repair of DSB, conjugational recombination, but also for the degradation of foreign linear DNA, and recently RecBCD was shown to participate in completion of DNA replication (45, 46). We unravel here that, in addition to these many functions, RecBCD also plays a role in SSG repair. Combining our original genetic system to monitor specifically HDGR events with different genetic backgrounds, we show that RecBCD plays a non-catalytic role in HDGR pathway. We also demonstrate that the RecBCD complex is necessary for efficient TLS bypass. Therefore, on the basis of our results, we propose that RecBCD plays a structural role, most likely by preserving the stability of the stalled replication fork to promote an efficient bypass of the lesion, either by TLS or HDGR pathway.

All previous known functions of RecBCD require its nuclease and helicase activities. However, by using specific alleles of RecB (*recB^D1080A^*) and RecD (*recD^K177Q^*), deficient respectively for the nuclease and helicase activities, we clearly demonstrate that they are dispensable for HDGR and TLS mechanisms. Our *in vivo* data that show that gap-repair does not require RecBCD nuclease nor helicase activities is in good agreement with previous *in vitro* data showing that RecBCD cannot unwind a ssDNA gap, but requires a blunt or nearly blunt double stranded end, and that its nuclease activity is very weak on a gapped substrate (47).

Until now, RecBCD had never been associated with SSG repair unless the ssDNA gap was converted into a DSB (48), its preferred substrate. In our context, it is very unlikely that the SSG generated at the lesion site is converted into a DSB. Indeed, if this were the case, i) RecBCD would unwind and resect an extended region of DNA which would require both its nuclease and helicase activities when we actually show that these activities are not required for SSG repair; ii) such resection would lead to the loss of our genetics markers, while we show that the genetic markers remain stable in the presence of functional RecBCD (4). Further evidence indicating the absence of DSB at the DNA lesion site came from the analysis of the *recD* deficient strain. This strain has been shown to be still proficient in DNA recombination and DSB repair (36), however when we monitor the level of HDGR in this strain, we observe a decrease similar to the one observed in the *recB-* strain. These data point towards a different role of RecBCD in DSB and SSG repair: while RecBC are sufficient for DSB repair (with the help of the RecFOR pathway (20), all components of the RecBCD complex are necessary for SSG repair. The *recB^D1080A^* mutant, whose nuclease is inactivated, conserves its helicase activities and is therefore still proficient for DSB repair provided that the RecFOR complex is present to load RecA. In the absence of RecFOR however, *recB^D1080A^* becomes deficient in DSB repair. By combining this allele with the *recF* or *recO* gene deletion, we observed no effect neither on HDGR nor on TLS compared to the *recF* or *recO* single mutant. This again rules out the possibility of RecBCD acting through the repair of a DSB.

After excluding a possible role of the helicase and nuclease domains of RecBCD, we hypothesised that the functional role of RecBCD in HDGR pathway could be to load RecA onto the single strand gap, together with (or in support to) the RecFOR complex, the other known mediator. This could explain the strong phenotype (*i.e*. similar to a *recA* deficient strain) we observed in the absence of both RecF and RecB (4). However, when using the *recA730* allele, that can load itself onto ssDNA without the help of its mediators, we could not complement the deficiency in HDGR in the *recB-* strain, while *recA730* allele could partially complement the defect in a *recF-* strain. This confirms that i) the decrease in HDGR observed in a *recF-* strain is indeed due to a defect in RecA loading and ii) RecBCD is not involved in mediating the loading of RecA to a ssDNA gap. This observation is also corroborated by the analysis of the *recB^D1080A^* strain. During DSB repair, the mutation in the nuclease domain has been shown to affect the RecA loading activity and the RecB^D1080A^CD complex was then dependent on the mediator activity of RecFOR complex (31, 33). In our lesion tolerance assay however, the double mutants *recF- recB^D1080A^* and *recO- recB^D1080A^* show a level of HDGR similar to the single *recF-* and *recO-* mutants indicating that RecBCD is not an alternative mediator of RecA in SSG repair.

Since the nuclease and helicase activities of RecBCD do not participate in HDGR, and no RecA loading activity was evinced, we surmise that RecBCD functions as a structural element in the HDGR pathway. One possible structural role could be the stabilization of the stalled replication fork. It is important to preserve the integrity of a stalled replication fork to avoid fork collapse, which in turn can lead to DSB formation that can be lethal for the cell. If such stabilization of the replication fork would occur, we reasoned that it would favor not only HDGR, but would also affect TLS. In *E. coli* under non-stressed conditions, TLS events represent a minor pathway compared to the HDGR pathway (22, 49). Since the basal level of TLS at the TT(6-4) lesion is very low (<0.5%), it is difficult to observe a clear decrease in TLS following the inactivation of the *recB* gene (Figure 5). The guanine adducts G-AAF and dG-BaP(-) are more frequently bypassed by TLS polymerases (Pol II/ Pol V and Pol IV, respectively) and inactivation of the *recB* gene in the presence of one of these two lesions results in a substantial decrease of TLS mediated by all three TLS polymerases. This result indicates that RecBCD complex plays a role not only in the HDGR pathway but also in the TLS pathway. The effect on the TLS pathway is not dependent on the catalytic activities of RecBCD neither on the repair of a DSB since we do not observe a decrease of TLS in the nuclease and helicase dead mutants nor in the DSB-repair deficient strains (*recF- recB^D1080A^* and *recO- recB^D1080A^*). Altogether these data support the hypothesis that RecBCD plays a structural role in SSG repair, allowing an efficient filling of the gap by HDGR or by TLS. This structural function seems to be more important for TLS, since the absence of RecBCD can lead to a decrease of up to ∼80% in TLS events (for dG-BaP), whereas it leads to a decrease of only ∼20% in HDGR events. It may be that the way the RecBCD complex participates to DDT is different for TLS and HDGR. Indeed, we observed a slight but reproducible difference of HDGR events between the leading and lagging strand for *recB* and *recD* mutants. The defect in HDGR is stronger when the lesion is located on the lagging strand for the *recB* strain, whereas it is stronger on the leading strand for the *recD* strain. However, no significant strand bias was observed for TLS in the *recB* strain. This leading/lagging strand difference for the *recB* and *recD* mutants could be explained by the opposite polarity of the helicases harbored by these two subunits. RecB possesses a 3′5′ helicase whereas RecD possesses a 5′3′ helicase. Since the inactivation of the helicase activity of RecD (*recD^K177Q^* strain) didn′t affect the DDT pathways, we rule out the involvement of this catalytic activity. However, it is possible that the protection of the replication fork occurs through the affinity of the helicases for DNA (independently of their activity). Following this hypothesis, the polarity of the RecB helicase would favor binding to the lagging strand whereas RecD would favor protection of the leading strand.

Our finding suggests that the RecBCD complex plays a structural role in SSG repair, most likely preserving the integrity of the stalled replication fork, which is important for both the TLS and HDGR pathways. However, how RecBCD does that still needs to be clarified. We propose that RecBCD binds or somehow protects the 3’-end of dsDNA-ssDNA junction of the stalled replication fork that can be the target of nucleolytic degradation operated by specific nucleases. Degradation of the nascent strand can be detrimental for the activity of the TLS polymerases, since the 3’-end at the lesion site is their cognate substrate. This would explain why inactivation of the *recB* gene alone has a stronger impact on TLS than on HDGR. If the extent of the degradation is not controlled and becomes too important, this will most likely also affect HDGR. If another lesion is present on the opposite strand, an extended resection of the non-protected 3′-end would eliminate the ds-DNA substrate necessary for homologous recombination at the lesion. The incapacity to repair the gap at either lesion will lead to cell death. This is indeed what we observe in the *recB*- and *recD*- strain where the decrease in HDGR is accompanied by a concomitant decrease in survival rather than an increase in damaged chromatid loss. This situation is different than when the RecFOR complex is absent: in that case, HDGR is affected by the delay in RecA loading at the gap, but the 3′-end is not resected and homologous recombination can occur if another lesion is present in the opposite strand.

In the last few years, a similar structural role has been proposed for the breast cancer susceptibility gene 2 (BRCA2) in human cells (50). BRCA2 is a key factor in homologous recombination during DSB repair where it recruits Rad51 (the functional homolog of RecA) to the ssDNA, but it is also involved in other DNA repair processes (reviewed in (51)). Several lines of evidence suggest that BRCA2 protects the stalled replication fork from undesired and harmful nucleolytic degradation, however the underlying molecular mechanisms still need to be completely elucidated (50, 52).

In conclusion, we show here that RecBCD plays a non-catalytic role in SSG repair, seemingly by preserving the integrity of the fork and allowing an efficient bypass of the lesion by both TLS and HDGR.

## ACKNOWLEDGEMENTS

We thank Élodie Chrabaszcz and Jean-Hugues Guervilly for helpful experimental discussions. We thank Pierre-Henri Gaillard and Mauro Modesti for critical reading of the manuscript. We thank Nick Geacintov for the dG-BaP (-) modified oligonucleotides.

## FUNDING

This work was supported by the Agence Nationale de la Recherche (ANR) grant GenoBlock [ANR-14- CE09-0010-01]. Funding for open access: ANR.

